# String-pulling by the common marmoset

**DOI:** 10.1101/2024.10.25.620352

**Authors:** Mathilde Bertrand, Michael Karkuszewski, Rhonda Kersten, Jean-Jacques Orban de Xivry, J. Andrew Pruszynski

## Abstract

Coordinated hand movements used to grasp and manipulate objects are crucial for many daily activities, such as tying shoelaces or opening jars. Recently, the string-pulling task – which involves cyclically reaching, grasping and pulling a string – has been used to study coordinated hand movements in rodents and humans. Here we characterize how adult common marmosets perform the string-pulling task and describe changes in performance across the lifespan. Marmosets (n=15, 7 female) performed a string-pulling task for a food reward using an instrumented apparatus attached to their home-cage. Movement kinematics were acquired using markerless video tracking and we assessed individual hand movements and bimanual coordination using standard metrics. Marmosets oriented their gaze towards the string above their hands and readily performed the task regardless of sex or age. The task required very little training and animals routinely engaged in multiple pulling trials per session despite not being under water or food control. All marmosets showed consistent pulling speed and similar hand movements regardless of age. Adult marmosets exhibited a clear hand effect, performing straighter and faster movements with their right hand despite showing idiosyncratic hand preference according to a traditional food retrieval assay. Hand effects were also evident for younger animals but seemed attenuated in the older animals. In terms of bimanual coordination, all adult marmosets demonstrated an alternating movement pattern for vertical hand positions. Two younger and two older marmosets exhibited idiosyncratic coordination patterns even after substantial experience. In general, younger and older animals exhibited higher variability in bimanual coordination than did adults.

## Introduction

Many real-world skills – tying a shoelace, buttoning a shirt, opening a jar – require performing skilled movements with each hand and coordinating the hands towards a common goal^1^. Understanding the neural basis of skilled hand movements^2–4^ and their bimanual coordination^5^ is of central interest in biological motor control, with the bulk of the animal work on this topic focused on dexterous non-human primates, especially the macaque monkey. Despite being less dexterous, recent work in the marmoset monkey demonstrates that they produce sophisticated reaching actions during prey capture in the wild^6^. When capturing moving crickets in a naturalistic laboratory setting, marmosets use a power grasp as opposed to a key or precision grip, but their grasp aperture scales with reach velocity and their movements appear to reflect predictive control strategies that compensate for sensorimotor delays^7^. These results, along with the many behavioral and anatomical similarities between marmosets and more dexterous primates^8–10^, suggest that the marmoset may serve as a suitable animal model for studying some aspects of reaching and grasping, which would be attractive given the many practical and technical advantages also afforded by the marmoset model^11–13^.

Here we further this effort by documenting marmoset behavior during unrestrained string-pulling, an ecologically valid behavior that includes both skilled hand movements and bimanual coordination. String-pulling involves making hand-over-hand movements to reel in the string or to acquire a reward attached to the string. It is an established paradigm in comparative psychology, with variations of the task being applied to over 100 species mostly to assess cognitive functions such as means-ends understanding^14^. Recently, string-pulling has been used to examine and compare motor control in rodents and humans^15–18^. Rodents and humans share many topographic and kinematic features when string-pulling, the most striking difference being that rodents rely on tactile cues from the vibrissa to guide their hands to the string whereas humans use vision to do so. String-pulling may also serve as a clinical biomarker since kinematic metrics appear sensitive and reliable enough to detect movement changes following neurological damage and muscle injury both in humans and animal models^17,19–23^.

We show that marmosets perform the string-pulling task in their group housed home-cage environment without extensive learning, naturally exhibiting cyclical alternating hand movements consistent with previous rodent and human studies^15,16,18^. Like humans and unlike rodents, marmosets look up at the string above their hands and do not touch it with their face; marmosets also appear to pre-shape their hand towards the appropriate grasping configuration in advance of string contact. Notably, our data indicate that, despite showing idiosyncratic hand preferences when retrieving food as has been previously reported^24,25^, adult marmosets produce straighter and faster reaches with their right hand when string-pulling, indicating some degree of motor lateralization. Lastly, we report that individual hand movements are relatively similar between adult, younger and older marmosets, but that older animals may show weaker lateralization and that both younger and older animals seem to exhibit higher variability in bimanual coordination.

## Methods

### Subjects

This study was conducted with fifteen common marmosets (*Callithrix jacchus*), 8 males and 7 females, from 6 to 168 months old (**Table 1**). Eight adults (4 males, 4 females; 25-60 months) were used for detailed characterization of string-pulling behavior. To qualitatively investigate age-related changes, animals were divided into three age groups: younger (<24 months, n=4); adult (as described above); and older (>90 months, n=3)^26^. All marmosets were housed in pairs or groups in a temperature (25 ± 1°C) and humidity (55 ± 15%) controlled facility with a 12-hour light-dark cycle. Marmosets had unrestricted access to food and water. All procedures were approved by the Animal Care Committee of the University of Western Ontario.

**Table 1:** Marmoset characteristics and hand preferences.

| *Subject* | *Sex* | *Age*<br>*(months)* | *Hand*<br>*Preference* | *Handedness*<br>*Index* |
| --- | --- | --- | --- | --- |
| *Y1* | F | 6 | R | 0.62 |
| *Y2* | M | 12 | N/A | N/A |
| *Y3* | M | 15 | R | 0.66 |
| *Y4* | F | 20 | L | -0.97 |
| *F1* | F | 28 | R | 0.23 |
| *M1* | M | 35 | Ambi | 0.00 |
| *M2* | M | 38 | L | -1.00 |
| *F2* | F | 49 | L | -0.83 |
| *F3* | F | 53 | R | 0.27 |
| *M3* | M | 56 | R | 0.93 |
| *M4* | M | 59 | R | 0.97 |
| *F4* | F | 59 | L | -0.73 |
| *O1* | M | 110 | L | -0.53 |
| *O2* | F | 114 | L | -0.90 |
| *O3* | M | 168 | Ambi | 0.10 |

### Handedness index

Hand preference was assessed using a simple task where marmosets retrieved a reward (a piece of marshmallow) by reaching through a hole into a box attached to their home-cage. Each trial was scored as left or right based on the hand used. Trials in which both hands were used were excluded. Each marmoset completed 20 trials per session across three sessions, resulting in a total of 60 trials, except for one younger male marmoset (Y2), for whom handedness was not assessed. A handedness index was calculated by subtracting the number of left-handed responses from the number of right-handed responses, and dividing by the total number of responses^27^ (**Table 1**). As such, the handedness index ranged from −1.0 (left-hand bias) to 1.0 (right-hand bias). Marmosets were classified as left-, ambidextrous, or right-handed based on binomial z-scores, calculated from the frequency of left- and right-hand responses. A z-score of −1.64 or lower indicated left-handed marmosets, while a z-scores of 1.64 or higher indicated right-handed marmosets. Intermediate scores were classified as ambidextrous.

### String-pulling apparatus

Our string-pulling apparatus was based on a previous rodent study^17^, using a transparent rectangular box (28 cm x 28 cm x 20 cm) connected to the home-cage. A loop of nylon string (4 mm in diameter) was attached to a continuous four-pulley system, one of which was connected to a rotary encoder to measure string speed. The pulley system was suspended by an aluminum frame constructed from 30 x 30 mm aluminum extrusions (Zyltech). A high-speed camera (240 FPS, GoPro / HERO 8) was facing the animal and collected data for offline analysis. The location of the camera relative to the transparent box was fixed across sessions (**Fig. 1A**).

**Figure 1:**
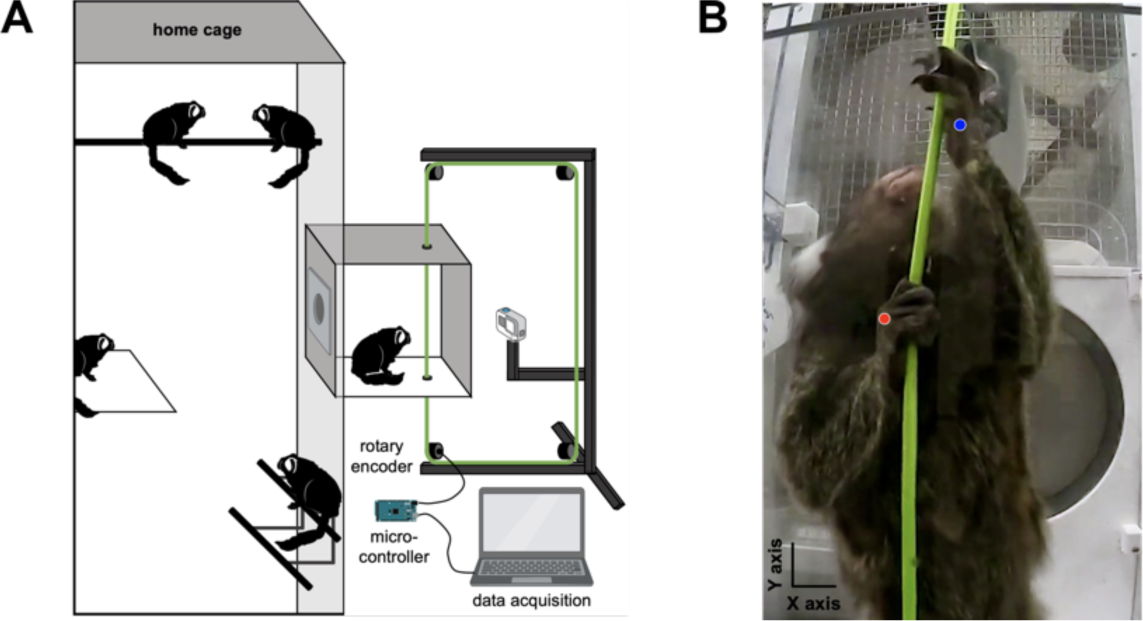
String-pulling apparatus. (**A**) Schematic of the string-pulling apparatus connected to the home-cage. (**B**) Frame from the string-pulling task. Note the upward gaze towards the reward. Red and blue dots represent the right- and left-hand positions according to the motion tracking algorithm.

The string-pulling task, in which animals had to pull a string to obtain a reward (a piece of marshmallow), was adapted from previous rodent and human studies^15,16,18,28^. During one-week of habituation, all marmosets spontaneously pulled on the string, then data were collected over five days. Only trials where the animal was facing the camera were included, and sessions with less than five trials were excluded. String speed was measured using a two-phase rotary encoder with a resolution of 600 pulses per rotation. Signals from the encoder were processed by a microcontroller (Arduino UNO Rev3, Arduino) to calculate rotation speed, direction (up or down), and length of string pulled. Initial calibration of the number of pulses per millimeter of string pulled was determined by manually pulling the string over a known distance.

### Behavioral analysis

A trial was defined as pulling the string until obtaining an attached reward, about 35 cm directly above the animal such that it could see the reward. At least 25 trials per animal were included over 5-day sessions. Movement topography and kinematics were averaged across all trials. All post-processing and filtering were performed using Matlab (R2022b, The MathWorks Inc., Natick MA, USA) and Python (version 3.9). Average string speed was extracted from the rotary encoder. Video recordings were trimmed with *ffmpeg* v4.4.2 to contain only the string-pulling behavior. Markerless motion tracking was conducted with DeepLabCut (v2.2.0)^29^, using a ResNet50 deep convolutional neural network model to extract locations and labels of the nose, string, right hand, and left hand (**Fig. 1B**). The known height of the apparatus was used to calculate pixel-to-centimeter conversion. The resulting traces were Butterworth filtered (0.75-9 Hz bandpass).

String-pulling involves a repeated, organized hand-over-hand sequence, consisting of a reaching phase, defined as upward forelimb movement without string contact, and a withdrawing phase, defined as downward movement that includes string contact before releasing the string. In addition, string-pulling cycles were analyzed manually, frame by frame, to identify six segments based on hand shape and location, as described in previous studies in humans and rodents^15–18^: lift (raising the hand), advance (slight adduction of the hand toward the string), grasp (flexing the fingers to grasp the string), pull (lowering the string by flexing the lower arm), push (lowering the string by extending the lower arm), and release (extending the fingers to release the string). Like rodent studies, we calculated the correlation between the nose and the string position to see if marmosets used somatosensory inputs from their face to track the string^15,17^.

Based on previous studies^15–17,21^, we calculated the most commonly used metrics associated with single hand movement during these phases: temporal topography (X and Y coordinates of each hand over time); heading direction (transformation of the origin coordinates (0,0) and the angle of the end coordinates of each hand relative to a polar coordinate system, when 0° is right, 90° is up, 180° is left, and 270° is down); heading variance (circular variance, i.e., concentration of the heading direction points); Euclidean distance (shortest distance between the start and end of the movement); path circuitry (ratio of the Euclidean distance to actual movement displacement, from zero, most circuitous path, to one, direct straight line); movement scaling (correlation between the set of peak speeds and Euclidean distances); peak speed (maximum speed in cm/s); and average speed (cm/s). Similarly, we calculated commonly used metrics associated with bimanual coordination: hand correlation (horizontal -X and vertical -Y correlation between both hands); and phase shift (temporal displacement of the waveforms of the X and Y hand coordinates).

### Statistical analysis

Statistical analyses were performed using GraphPad Prism 8 (GraphPad Software Inc., USA). For single-hand metrics in adult animals, three-way ANOVAs were used to analyze the effect of hand, phase and sex. For bimanual coordination metrics, Wilcoxon signed-rank tests were used to compare each median with zero. Pearson correlations were calculated to investigate the association between age and string speed. Differences were considered statistically significant if p<0.05.

## Results

The string-pulling apparatus was mounted to the home-cage of a group housed set of marmosets (**Fig. 1**). After a short habituation period, marmosets willingly entered the apparatus. Only one marmoset was allowed to enter the apparatus at a given time. Access to the apparatus was controlled by the experimenter via a sliding door at the entrance. Once inside, the marmoset remained in the apparatus until it showed no further interest in performing the task, at which point the sliding door was opened and it could return to its home-cage. String-pulling was always done in full-light conditions and no attempts were made to alter the normal housing setup or the arrangement of the room, which included multiple homes cages.

### String pulling behavior in adult marmosets

All eight adult marmosets spontaneously engaged in the string-pulling task, often from their very first exposure. Adult marmosets seemed to perform the task guided by vision. Like humans^18^, marmosets appeared to keep their gaze on the string above the grasp point (**Fig. 1B**, **Fig. 2A**). We found no correlation between marmoset nose position and string position in the horizontal direction (median correlation across animals: r = −0.008, Wilcoxon signed-rank test against zero: p=0.13). That is, like humans and unlike rodents^15,16^, marmosets did not appear to use somatosensory inputs from their face to guide the string.

**Figure 2:**
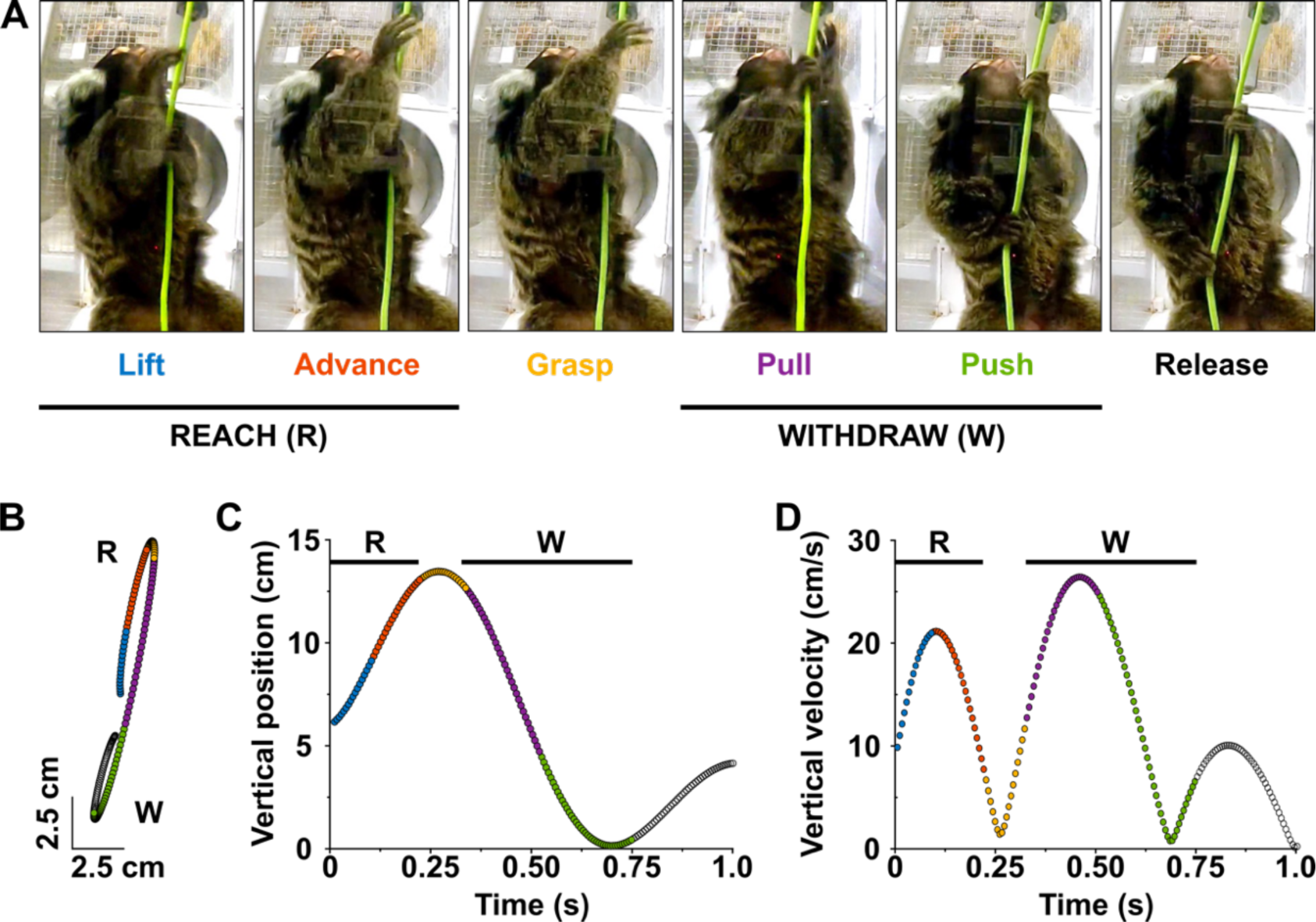
Example of a single string-pulling cycle for the right hand. (**A**) Sequence of six movement segments during string pulling: *Lift* (blue), the hand is raised; *Advance* (orange), hand is extended towards the string; *Grasp* (yellow), fingers are flexed to grasp the string; *Pull* (purple), the string is lowered by flexing the lower arm; *Push* (green), the string is lowered by extending the lower arm; *Release* (black), fingers are opened and extended to release the string. The reach phase (R) is defined as upward movements without string contact (lift and advance), the withdraw phase (W) is defined as downward movements with string contact (pull and push). Note the gaze directed on the string, above the grasp point. (**B**) Spatial segmentation of the six segments. Dots are plotted every 6 ms. (**C**) Vertical position of the segments over time. Data is aligned on the beginning of Lift. (**D**) Speed of the segments over time.

In terms of individual hand movements, string-pulling in adult marmosets was qualitatively similar to what has previously been described in mice, rats, and humans. Marmosets showed the typical reach-and-withdraw cycle, with a reach phase involving upward movements to grasp the string without string contact, and a withdraw phase involving downward movements while grasping the string before releasing it. Marmoset grasp aperture opened during the reaching phase and generally started to close before contact was made with the string (**Fig. 1B, 2A**).

We divided each reach cycle into six segments based on hand position and orientation: lift (raising the hand), advance (slight adduction of the hand toward the string), grasp (flexing the fingers to grasp the string), pull (lowering the string by flexing the lower arm), push (lowering the string by extending the lower arm), and release (extending the fingers to release the string) (**Fig. 2A**). Typically, hand speed increased before grasping the string, peak reach speed occurred near the transition between the lift and advance phases, approached zero during the grasp phase, increased again during the push and pull phase of the withdrawal and approached zero during release (**Fig. 2B-D**).

We characterized the reach and withdraw phases using several standard metrics (see Methods). First, we calculated the heading of the movement trajectories (**Fig. 3A**). In addition to moving in opposite direction for the reach and withdraw phases (as required by the task), marmosets moved their hands across their body for both phases, yielding a reliable difference in heading direction between hands (main effect of hand: F_1,12_=19.63, p=0.0008), which did not differ between males and females (main effect of sex: F_1,12_=0.064, p=0.80).

**Figure 3:**
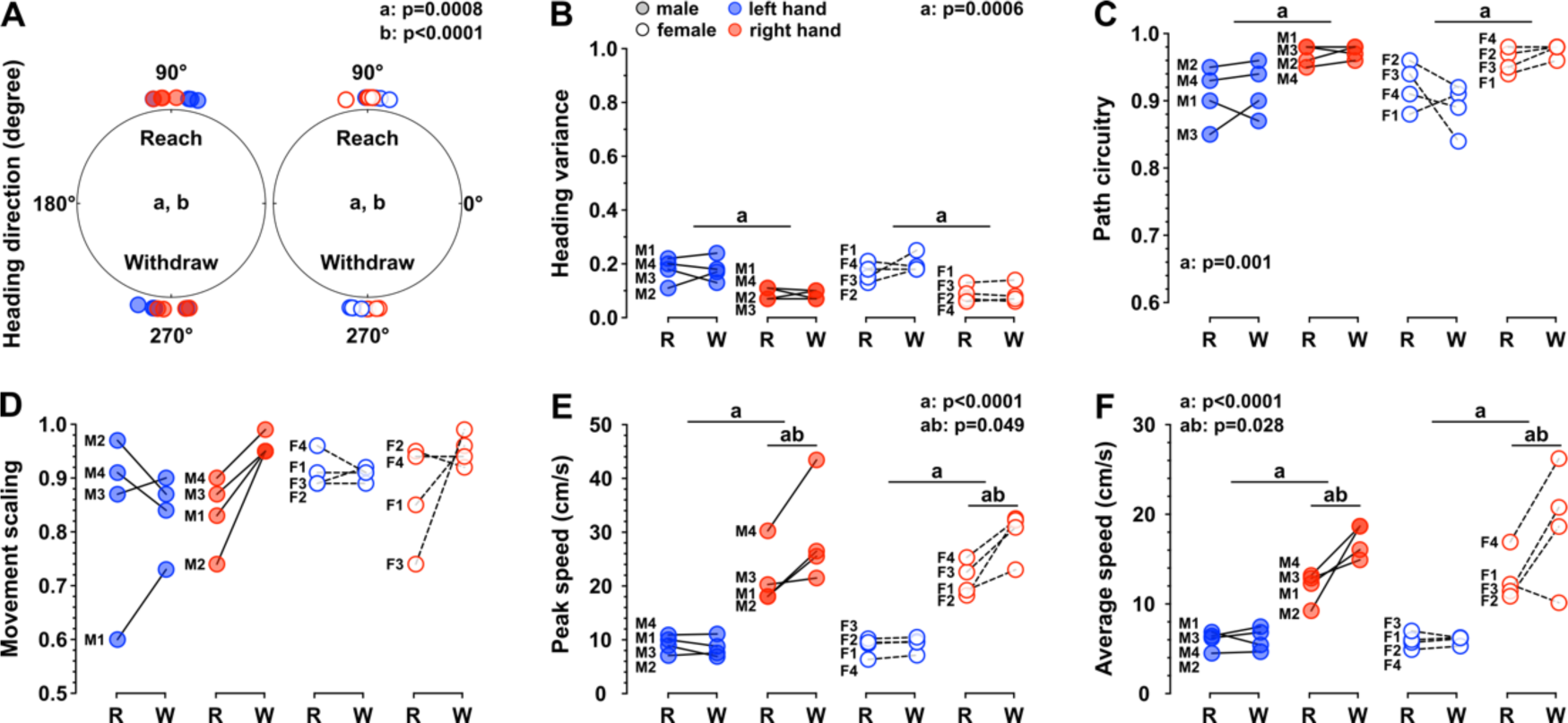
Single hand metrics. Metrics for the left (blue) and right (red) hand during the reach (R) and withdraw (W) phases. Medians of (**A**) heading direction, (**B**) heading variance, (**C**) path circuitry, (**D**) movement scaling, (**E**) peak speed, and (**F**) average speed are shown for each adult male (M) and female (F) marmoset. Main effect of three-way ANOVA: a, hand; b, phase; ab, hand by phase interaction.

Second, we examined movement consistency by calculating the variance of heading trajectory across reaches (**Fig. 3B**). Marmosets were very consistent, exhibiting a low variance, and even more consistent with the right hand than the left hand (main effect of hand: F_1,12_=21.38, p=0.0006). This consistency was similar across phases (main effect of phase: F_1,12_=0.35, p=0.57) and as a function of sex (F_1,12_=0.66, p=0.43).

Third, we calculated path circuitry, a measure of movement straightness (**Fig. 3C**). Although adult marmosets showed relatively straight movement trajectories, right hand trajectories were consistently straighter than left hand trajectories (main effect of hand: F_1,12_=17.81, p=0.001). Straightness was not reliably different between reach and withdraw phases (main effect of phase: F_1,12_=1.41, p=0.26), or between males and females (main effect of sex: F_1,12_=0.56, p=0.47).

Fourth, we calculated movement scaling, the correlation between peak speed and the Euclidean distance of each movement (**Fig. 3D**). All marmosets scaled their movements similarly (main effect of sex: F_1,12_=2.26, p=1.16), with both hands (main effect of hand: F_1,12_=1.42, p=0.26) and for both phases (main effect of phase: F_1,12_=3.64, p=0.08).

Fifth, we calculated peak and average pulling speed (**Fig. 3E,F**). All adult marmosets pulled faster with their right hand (main effect of hand: peak speed: F_1,12_=80.37, p<0.0001; average speed: F_1,12_=68.14, p<0.001) and even somewhat faster during the withdraw phase for their right hand (hand by phase interaction: peak speed: F_1,12_=4.79, p=0.049; average speed: F_1,12_=6.22, p=0.03). Like the other metrics, we observed no reliable difference between males and females (main effect of sex: peak speed: F_1,12_=0.004, p=0.95; average speed: F_1,12_=0.58, p=0.46).

We characterized bimanual coordination by examining the relationship between horizontal and vertical trajectories across the hands. We found in-phase horizontal movements between hands (**Fig. 4A, C, E**), presumably reflecting the general need to move the hands together to account for the shifting horizontal position of string over time. We also found largely anti-phasic vertical movements (**Fig. 4B, D, F**), representing the alternating movements that define the cyclical string-pulling action.

**Figure 4:**
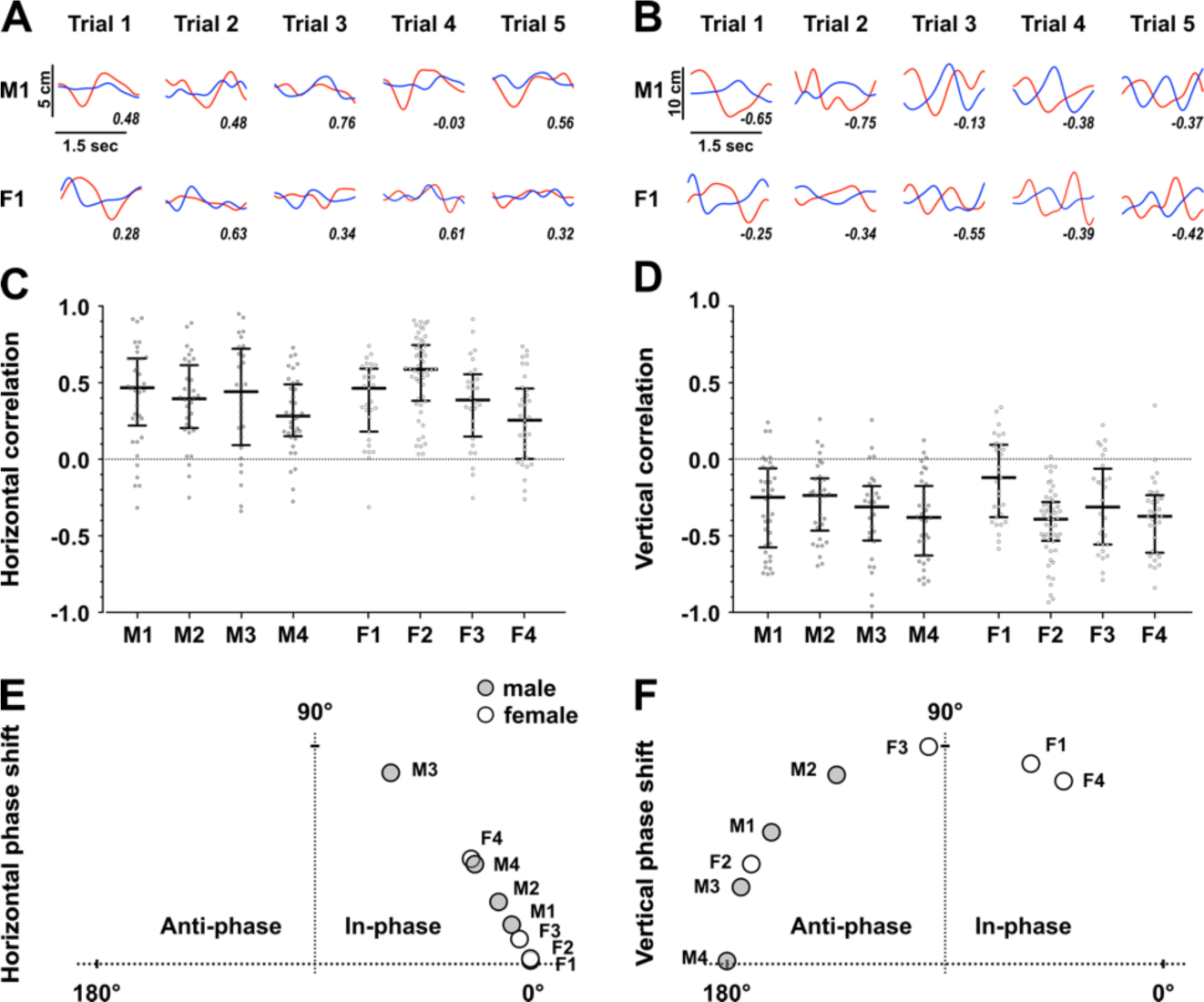
Bimanual coordination. Example of left (blue) and right (red) hand position on the horizontal X-axis (**A**) and vertical Y-axis (**B**) for five trials of a male (M1) and a female (F2) marmoset, with corresponding correlation values. Medians of horizontal (**C**) and vertical (**D**) correlations between left and right hands, with data points for each trial for each male (M) and each female (F) marmosets. Median of horizontal (**E**) and vertical (**F**) circular phase shift representing the in-phase and anti-phase movements between left and right hands for each marmoset.

### String-pulling across the lifespan

We examined string-pulling behavior in a small number of younger (<24 months old) and older marmosets (>90 months old) in our colony. The main goal of examining these groups was to establish whether they could perform the task as motivation for future developmental and aging studies. Given the small sample size we did not perform the same statistical analysis as for the adults but present the results in a similar fashion for qualitative comparison.

Like the adults, both younger and older marmosets spontaneously performed the string-pulling task. Although all animals readily learned the task, it took more training days to obtain five sessions for older marmosets because they were less likely to produce the number of required string-pulling trials early in training. The limiting factor with respect to learning appeared to be motivation rather than movement control. After habituation, we found no reliable correlation between age and average string speed (Pearson r=0.35, p=0.20; **Fig 5**). Even our oldest animal pulled faster than some adult animals.

**Figure 5:**
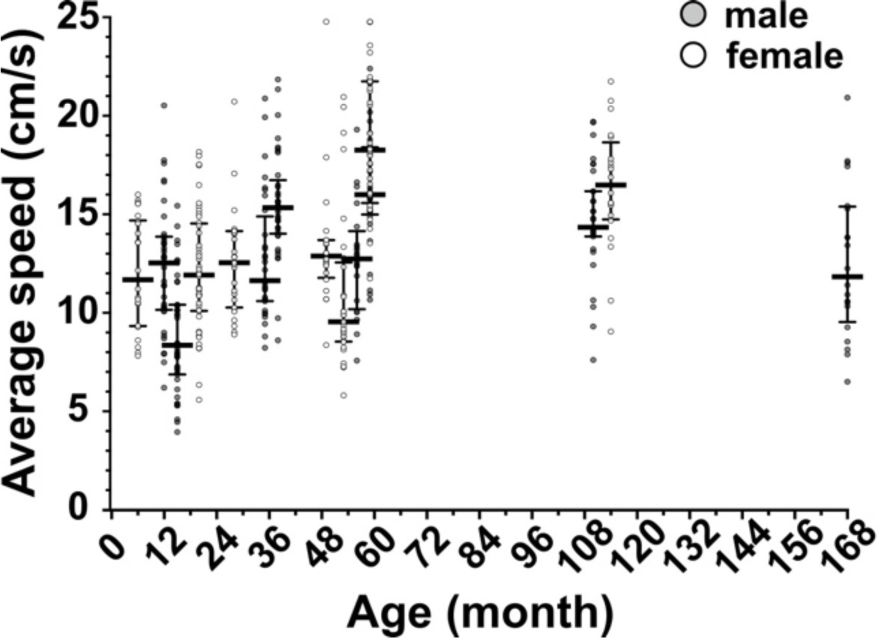
String pulling speed across the lifespan. String speed as extracted from a rotary encoder for each marmoset. The individual dots represent single trials. The horizontal line represents the median for each animal.

Using the same single-hand metrics as for the adult marmosets above, we quantified the reach and withdrawal phases of each hand for younger and older marmosets, which generally showed the same trends as the adult animals (**Fig. 6**). Younger and older animals appeared to make hand movements across their body for both reach and withdrawal phases **(Fig. 6A)**. Both groups appeared to exhibit greater movement consistency for the right hand than the left hand, though this lateralization might have been reduced in the older animals (**Fig. 6B**). Reduced lateralization was also apparent in movement straightness (**Fig. 6C**). On the other hand, similar to the adults, younger and older animals appeared to show a marked increase movement scaling for withdrawal movements with their right hand (**Fig. 6D**) and increased movement speed with the right hand in general (**Fig. 6E,F**).

**Figure 6:**
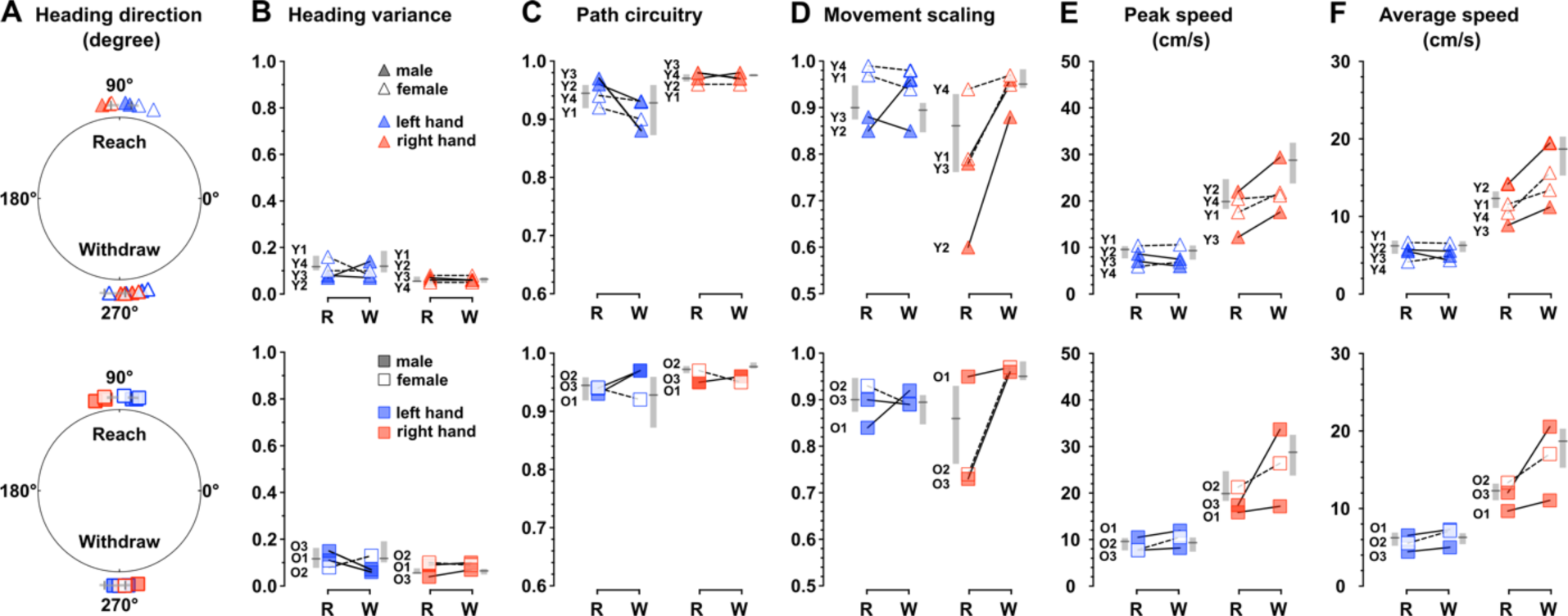
Single hand metrics across the lifespan. Left (red) and right (blue) hand metrics for younger (Y, <24 months) and older (O, >90 months, square) marmosets during the reach (R) and withdraw (W) phases. Medians of (**A**) heading direction, (**B**) heading variance, (**C**) path circuitry, (**D**) movement scaling, (**E**) peak speed, and (**F**) average speed are shown for younger (triangle, top panel) and older (square, bottom panel) marmosets. Grey lines and bars on the side of each metric represent median values and interquartile range for adult marmosets.

With respect to bimanual coordination, both younger and older marmosets displayed the typical cyclical string-pulling behavior, generally showing in-phase horizontal movements (**Fig. 7A, C**) and anti-phasic vertical movements between hands (**Fig. 7B, D)**. However, some of these animals also exhibited more variable and idiosyncratic coordination patterns, with no significant positive correlations in horizontal movements (Y2: p=0.27) (**Fig. 7C**), and/or no significant negative correlations in vertical movements (Y1: p=0.26; O1: p=0.37; O3: p=0.09) (**Fig. 7D**).

**Figure 7:**
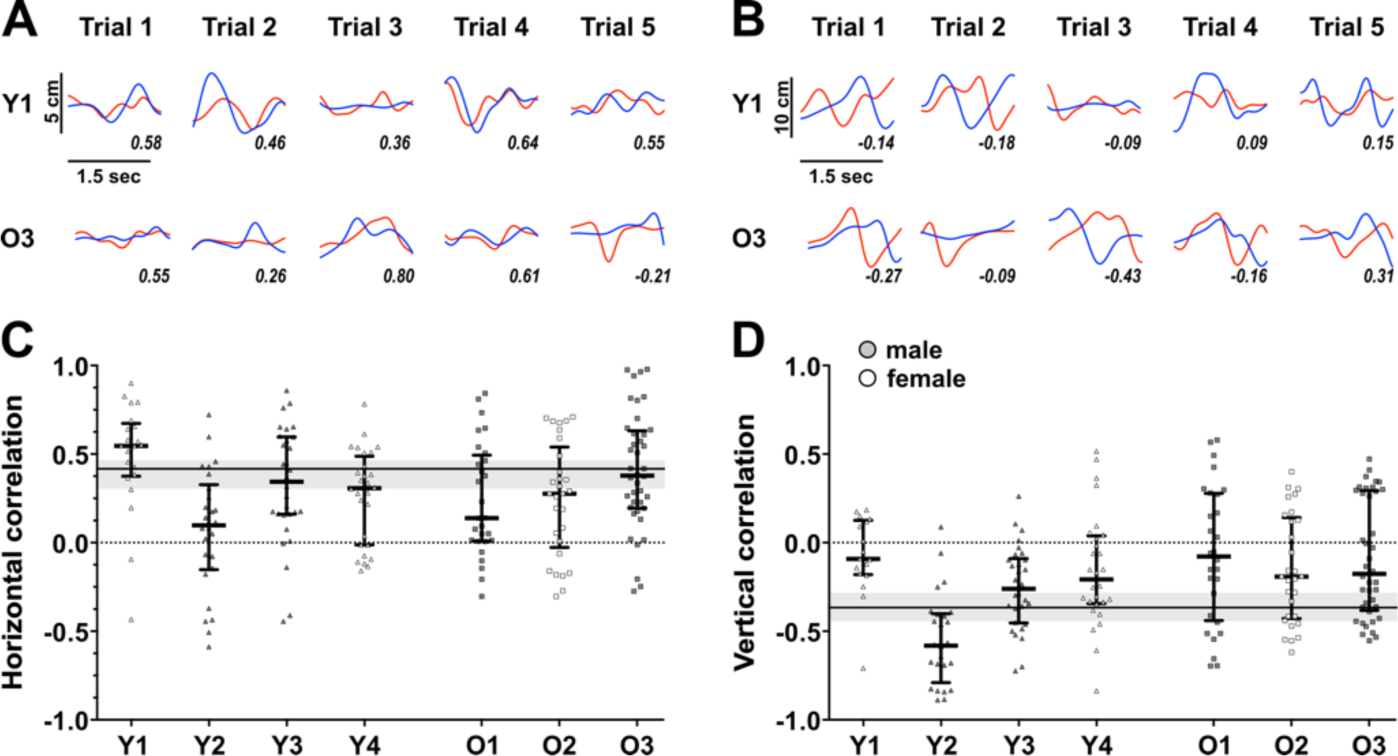
Bimanual coordination across the lifespan. Example of left (blue) and right (red) hand position on the horizontal X-axis (**A**) and vertical Y-axis (**B**) for five trials of one younger (Y1) and one older (O3) marmoset, with corresponding correlation values. Medians with interquartile range of horizontal (**C**) and vertical (**D**) correlations between both hands, with data points for each trial for each younger (Y, >24 months, triangle) and older (O, >90 months, square) marmoset. Black line and gray bar represent median and interquartile range from adult marmosets.

## Discussion

Our study provides the first characterization of string-pulling in the common marmoset. We show that marmosets, like rodents and humans, readily perform the string-pulling task with minimal training and naturally exhibit a cyclical alternating pattern of hand movements^15–18^. Like humans, but unlike rodents, marmoset string-pulling appears to be driven by vision^15,16,18^ and the animals seem to pre-shape their hand in advance of string contact, a pattern also observed when marmosets rapidly reach to capture moving prey^7^. Notably, despite showing idiosyncratic hand preferences when reaching for food, adult marmosets show a clear difference in their individual hand metrics during string-pulling, producing straighter, faster and less variable kinematics with their right hand. Our findings suggest some age-related changes during string-pulling. Although the overall behavior is quite similar between adult, younger and older marmosets, both younger and older animals demonstrate higher variability with respect to bimanual coordination. Older marmosets may have reduced hand effects with respect individual hand metrics.

### The string-pulling task for marmoset motor neuroscience

The string-pulling task has several key features, over and above the fact that it includes both skilled hand movements and bimanual coordination, which make it an attractive task for studying the neural control of movement in the marmoset monkey. String-pulling, like cricket catching^7^, is a well-controlled task that requires little to no training, presumably because it mimics ecologically conserved behaviors like climbing or foraging^14,30^. Indeed, male and female marmosets of all ages spontaneously engaged in the task within a week of first exposure even though they are under no water or food restriction. The ecological validity of the task may explain the relative similarity of the behavior across the lifespan compared to previous reports using tasks with higher cognitive and motor demands^31–34^. Although the basic string-pulling task may be an advantage when investigating some motor changes with aging, more complex or difficult variants may be needed if the goal is to reveal the fuller spectrum of age-related decline.

String-pulling affords many manipulations that are of direct relevance to sensorimotor neuroscience. For example, manipulating string thickness provides a way of studying the control of grasp aperture^35^ and online planning^36,37^. Changing the tension on the string or the texture of the string provide a way of studying the coordination of grip and load forces^38,39^, inducing motor learning^40^, and/or probing fast feedback responses to visual or mechanical perturbations^41–43^. Such manipulations are powerful because they require different control strategies, and since they do not change the natural string-pulling behavior, they can likely be enacted quickly if not in real-time, which is not generally the case in more abstract experimental settings.

As demonstrated here, string-pulling is readily amenable for in-cage studies which makes it possible to study unconstrained behavior but also to assess relatively large cohorts of marmosets, as is often needed for studying disease models and testing clinical interventions. At the same time, our ongoing work indicates that chair-restrained and head-fixed marmosets perform the string-pulling behavior in a similar manner. Future studies in this more controlled setting and with a multi-camera tracking setup will open up more precise experiments and a host of more sophisticated methods including large-scale electrophysiology via acutely inserted Neuropixel electrodes^44^ or Myomatrix arrays^45^, as well as calcium imaging^46^ and optogenetic stimulation^47^.

### String-pulling movements

All marmosets appeared to use vision to track the string, but they directed their gaze above their hands and generally did not look at the grasp point. This behavior is similar to human string-pulling, where participants focus their gaze at the grasp point only when they were asked to reach for a specific cue and otherwise look up at the string above their hands^18^. The observation that marmosets, like humans do not fixate on the grasp point, indicates that they rely on somatosensory inputs – both the tactile events that occur when string contact is made and proprioceptive signals about joint configuration – to guide their hands to the string. Mice and rats must also use somatosensory inputs to guide their hands since they grasp the string in an unseen location below their snout^15,16^. Interestingly, unlike marmosets and humans, rodents keep their snout and vibrissae in constant contact with the string. These vibrissae inputs in rodents, like visual inputs in primates, may provide online tracking of string position.

Marmoset string-pulling behavior involved hand-over-hand cycling movements with distinct reach and withdraw phases. Related studies in rodents and humans have identified a similar six segment sequence within the reach and withdraw phases (lift; advance; grasp; pull; push; release) characterized by specific trajectories, arm movements and hand shapes^15–18^. However, our results suggest that grasping strategies differ across species. Rodents use an arpeggio grip, first contacting the string with extended fingers, then closing all fingers at once based on somatosensory feedback^15,16,48^. Humans, by contrast, use a whole-hand grip, fully opening all fingers before the grasp, contacting the string with the finger 5 (pinky) and 4 to guide it toward the palm while sequentially closing the digits from finger 5 to 1 (thumb) to hold the string^18^.

Previous studies have shown that marmosets typically use a power grasp^16,49,50^, pre-shaping the hand by moving all fingers simultaneously before object contact and closing them around the object and pressing it against their palm. However, whether they scale their grip according to the size of the target remains debated^7,50,51^. We observed a similar power grasp pattern but our results cannot address the issue of scaling directly as we did not manipulate string thickness, and our use of a single frontal camera limited our analysis to relatively gross whole arm effects.

Adult marmosets exhibited direct, scaled, and consistent movements, with faster movements during the withdraw phase than the reach phase. This may result from a different utilization of gravitational force during reach (against gravity) and withdraw (with gravity) movements^52–54^. Studies have shown longer acceleration during downward movements compared to upward movement of equivalent duration and amplitude^55,56^. Although our present setup did not allow for it, it would be relatively straightforward to test this idea by having marmosets perform the string-pulling behavior in a horizontal apparatus, where the effects of gravitational force are limited. Lastly, marmosets stabilize their posture by pulling the string during the withdraw phase, which likely contributes to faster movements compared to the reach phase, where their hands move freely.

### Marmoset handedness

Although we did not observe a systematic bias in animal posture, some caution should be taken when interpreting kinematic differences between hands since our setup is relatively unconstrained, and our analysis is based on video from a single front-facing camera that cannot resolve out of plane movements. That said, we found that adult marmosets showed straighter, faster, and less variable movements with their right hand. This systematic right-hand ‘advantage’ in terms of movement characteristics did not correlate with a standard hand preference assay^27^ which, consistent with most previous work^24,57–59^, showed that about half of the marmosets in our sample had a right hand preference. The disconnect between movement features and hand preference may partly reflect known postural effects on hand preference. Right hand use in some primates is increased for bipedal reaching compared to quadrupedal reaching^60^ and our string-pulling task was performed in bipedal posture whereas the food preference assay was generally performed from a quadrupedal posture^61^. Interestingly, no equivalent hand specific effects were reported in rodent string-pulling studies^15–17^, despite evidence of hand preferences and brain asymmetry^62–64^. In human string-pulling, not all parameters reported here were examined but asymmetrical hand movements were only observed when the task was performed from memory and these were not hand specific^18^.

The relationship between brain lateralization, where each hemisphere exerts preferential control over certain behavioral functions^65–67^, and handedness, which refers to the preferred hand for doing manual tasks^68,69^, has been largely studied in humans^70–73^. One prominent idea is that sensorimotor laterization is related to the functional specialization of each arm for different functions^74–79^. For example, right-handed patients with left hemisphere damage after a stroke show deficits in arm trajectory control and joint coordination, whereas right-handed patients with right hemisphere damage show deficits in movement accuracy^80^. Given this previous work, it may be that the hand effect we report stems from such hemispheric specialization. However, our findings indicate that the specialization that yields hand preference can be decoupled from the specialization that yields hand specific movement characteristics. How this relates to humans is hard to say given the very strong population bias towards right-hand preference compared to other animals and the relative paucity of studies on human left-handers. That said, the studies that exist in this context suggest that lateralization with respect to hand preference and hand movement characteristics are related in that left-handers show an inverted pattern of movement characteristics though this inversion is not as strong and not completely symmetrical^81,82^. Lastly, our data suggest that the hand effect on movement characteristics is somewhat weaker for the older marmosets. Such a result, if born out with a properly powered sample, would be consistent with previous human studies showing a decrease in hemispheric asymmetry in older adults^83–86^ and thus provide an experimental window into its underlying neural mechanisms.

## Contributions

All authors conceived the project and designed the study. M.B. and M.K. collected the data and performed the analyses. M.B. wrote the first draft of the paper. All authors discussed the results and edited the manuscript.

## Acknowledgements

This work was supported by a CIHR Project Grant to JAP (PJT-186177), the Azrieli Foundation (via COMPERE, the Collaboration on Motor Planning, Execution and Resilience), and the Canada First Research Excellence Fund (BrainsCAN). JAP received a salary award from the Canada Research Chairs program.

## References

1. Obhi SS. Bimanual Coordination: An Unbalanced Field of Research. Motor Control. 2004;8(2):111–120. doi:10.1123/mcj.8.2.111

2. Sobinov AR, Bensmaia SJ. The neural mechanisms of manual dexterity. Nat Rev Neurosci. 2021;22(12):741–757. doi:10.1038/s41583-021-00528-7

3. Scott SH. The computational and neural basis of voluntary motor control and planning. Trends Cogn Sci. 2012;16(11):541–549. doi:10.1016/j.tics.2012.09.008

4. Arber S, Costa RM. Networking brainstem and basal ganglia circuits for movement. Nat Rev Neurosci. 2022;23(6):342–360. doi:10.1038/s41583-022-00581-w

5. Swinnen SP, Wenderoth N. Two hands, one brain: cognitive neuroscience of bimanual skill. Trends Cogn Sci. 2004;8(1):18–25. doi:10.1016/j.tics.2003.10.017

6. Ngo V, Gorman JC, De La Fuente MF, Souto A, Schiel N, Miller CT. Active vision during prey capture in wild marmoset monkeys. Curr Biol. 2022;32(15):3423–3428.e3. doi:10.1016/j.cub.2022.06.028

7. Shaw L, Wang KH, Mitchell J. Fast prediction in marmoset reach-to-grasp movements for dynamic prey. Curr Biol. 2023;33(12):2557–2565.e4. doi:10.1016/j.cub.2023.05.032

8. Bakola S, Burman KJ, Rosa MGP. The cortical motor system of the marmoset monkey (Callithrix jacchus). Neurosci Res. 2015;93:72–81. doi:10.1016/j.neures.2014.11.003

9. Kaas JH. Comparative Functional Anatomy of Marmoset Brains. ILAR J. 2020;61(2-3):260–273. doi:10.1093/ilar/ilaa026

10. Mitchell JF, Leopold DA. The marmoset monkey as a model for visual neuroscience. Neurosci Res. 2015;93:20–46. doi:10.1016/j.neures.2015.01.008

11. Mansfield K. Marmoset Models Commonly Used in Biomedical Research. Comp Med. 2003;53(4).

12. Miller CT, Freiwald WA, Leopold DA, Mitchell JF, Silva AC, Wang X. Marmosets: A Neuroscientific Model of Human Social Behavior. Neuron. 2016;90(2):219–233. doi:10.1016/j.neuron.2016.03.018

13. Okano H, Hikishima K, Iriki A, Sasaki E. The common marmoset as a novel animal model system for biomedical and neuroscience research applications. Semin Fetal Neonatal Med. 2012;17(6):336–340. doi:10.1016/j.siny.2012.07.002

14. Jacobs IF, Osvath M. The string-pulling paradigm in comparative psychology. J Comp Psychol. 2015;129(2):89–120. doi:10.1037/a0038746

15. Blackwell AA, Banovetz MT, Qandeel, Whishaw IQ, Wallace DG. The structure of arm and hand movements in a spontaneous and food rewarded on-line string-pulling task by the mouse. Behav Brain Res. 2018;345:49–58. doi:10.1016/j.bbr.2018.02.025

16. Blackwell AA, Köppen JR, Whishaw IQ, Wallace DG. String-pulling for food by the rat: Assessment of movement, topography and kinematics of a bilaterally skilled forelimb act. Learn Motiv. 2018;61:63–73. doi:10.1016/j.lmot.2017.03.010

17. Jordan GA, Vishwanath A, Holguin G, et al. Automated System for Training and Assessing String-Pulling Behaviors in Rodents. Bioengineering; 2023. doi:10.1101/2023.07.02.547431

18. Singh S, Mandziak A, Barr K, et al. Human string-pulling with and without a string: movement, sensory control, and memory. Exp Brain Res. 2019;237(12):3431–3447. doi:10.1007/s00221-019-05684-y

19. Blackwell AA, Schell BD, Osterlund Oltmanns JR, et al. Skilled movement and posture deficits in rat string-pulling behavior following low dose space radiation (28Si) exposure. Behav Brain Res. 2021;400:113010. doi:10.1016/j.bbr.2020.113010

20. Blackwell AA, Tracz JA, Fesshaye AS, et al. Fine motor deficits exhibited in rat string-pulling behavior following exposure to sleep fragmentation and deep space radiation. Exp Brain Res. 2023;241(2):427–440. doi:10.1007/s00221-022-06527-z

21. Darevsky DM, Hu DA, Gomez FA, Davies MR, Liu X, Feeley BT. A Tool for Low-Cost, Quantitative Assessment of Shoulder Function Using Machine Learning. Orthopedics; 2023. doi:10.1101/2023.04.14.23288613

22. Hart M, Blackwell AA, Whishaw IQ, Wallace DG, Cheatwood JL. Impairments and compensation in string-pulling after middle cerebral artery occlusion in the rat. Behav Brain Res. 2023;450:114469. doi:10.1016/j.bbr.2023.114469

23. Khademullah CS, De Koninck Y. A novel assessment of fine-motor function reveals early hindlimb and detectable forelimb deficits in an experimental model of ALS. Sci Rep. 2022;12(1):17010. doi:10.1038/s41598-022-20333-1

24. Cordeiro De Sousa MB, Xavier NS, Alves Da Silva HP, Souza De Oliveira M, Yamamoto ME. Hand preference study in marmosets (Callithrix jacchus) using food reaching tests. Primates. 2001;42(1):57–66. doi:10.1007/BF02640689

25. Hook MA, Rogers LJ. Development of hand preferences in marmosets (Callithrix jacchus) and effects of aging. J Comp Psychol. 2000;114(3):263–271. doi:10.1037/0735-7036.114.3.263

26. Abbott DH, Barnett DK, Colman RJ, Yamamoto ME, Schultz-Darken NJ. Aspects of Common Marmoset Basic Biology and Life History Important for Biomedical Research. Comp Med. 2003;53(4).

27. Vaughan E, Le A, Casey M, Workman KP, Lacreuse A. Baseline cortisol levels and social behavior differ as a function of handedness in marmosets (*Callithrix jacchus*). Am J Primatol. 2019;81(9):e23057. doi:10.1002/ajp.23057

28. Conner JM, Bohannon A, Igarashi M, Taniguchi J, Baltar N, Azim E. Modulation of tactile feedback for the execution of dexterous movement. Science. 2021;374(6565):316–323. doi:10.1126/science.abh1123

29. Mathis A, Mamidanna P, Cury KM, et al. DeepLabCut: markerless pose estimation of user-defined body parts with deep learning. Nat Neurosci. 2018;21(9):1281–1289. doi:10.1038/s41593-018-0209-y

30. Ford SM, Porter LM, Davis LC, eds. The Smallest Anthropoids: The Marmoset/Callimico Radiation. Springer US; 2009. doi:10.1007/978-1-4419-0293-1

31. Callahan DM, Kent-Braun JA. Effect of old age on human skeletal muscle force-velocity and fatigue properties. J Appl Physiol. 2011;111(5):1345–1352. doi:10.1152/japplphysiol.00367.2011

32. Fielding RA, Vellas B, Evans WJ, et al. Sarcopenia: An Undiagnosed Condition in Older Adults. Current Consensus Definition: Prevalence, Etiology, and Consequences. International Working Group on Sarcopenia. J Am Med Dir Assoc. 2011;12(4):249–256. doi:10.1016/j.jamda.2011.01.003

33. Power GA, Dalton BH, Rice CL. Human neuromuscular structure and function in old age: A brief review. J Sport Health Sci. 2013;2(4):215–226. doi:10.1016/j.jshs.2013.07.001

34. Wishart LR, Lee TD, Murdoch JE, Hodges NJ. Effects of Aging on Automatic and Effortful Processes in Bimanual Coordination. J Gerontol B Psychol Sci Soc Sci. 2000;55(2):P85–P94. doi:10.1093/geronb/55.2.P85

35. Johansson RS, Flanagan JR. Coding and use of tactile signals from the fingertips in object manipulation tasks. Nat Rev Neurosci. 2009;10(5):345–359. doi:10.1038/nrn2621

36. Ariani G, Kordjazi N, Pruszynski JA, Diedrichsen J. The Planning Horizon for Movement Sequences. eneuro. 2021;8(2):ENEURO.0085-21.2021. doi:10.1523/ENEURO.0085-21.2021

37. Kashefi M, Reschechtko S, Ariani G, et al. Future movement plans interact in sequential arm movements. eLife. 2024;13:RP94485. doi:10.7554/eLife.94485

38. Westling G, Johansson RS. Factors influencing the force control during precision grip. Exp Brain Res. 1984;53(2). doi:10.1007/BF00238156

39. Flanagan JR, Tresilian J, Wing AM. Coupling of grip force and load force during arm movements with grasped objects. Neurosci Lett. 1993;152(1-2):53–56. doi:10.1016/0304-3940(93)90481-Y

40. Krakauer JW, Hadjiosif AM, Xu J, Wong AL, Haith AM. Motor Learning. In: Terjung R, ed. Comprehensive Physiology. 1st ed. Wiley; 2019:613–663. doi:10.1002/cphy.c170043

41. Scott SH, Cluff T, Lowrey CR, Takei T. Feedback control during voluntary motor actions. Curr Opin Neurobiol. 2015;33:85–94. doi:10.1016/j.conb.2015.03.006

42. Pruszynski JA. Primary motor cortex and fast feedback responses to mechanical perturbations: a primer on what we know now and some suggestions on what we should find out next. Front Integr Neurosci. 2014;8. doi:10.3389/fnint.2014.00072

43. Pruszynski JA, Scott SH. Optimal feedback control and the long-latency stretch response. Exp Brain Res. 2012;218(3):341–359. doi:10.1007/s00221-012-3041-8

44. Dotson NM, Davis ZW, Jendritza P, Reynolds JH. Acute Neuropixels Recordings in the Marmoset Monkey. Neuroscience; 2023. doi:10.1101/2023.12.14.571771

45. Chung B, Zia M, Thomas KA, et al. Myomatrix arrays for high-definition muscle recording. eLife. 2023;12:RP88551. doi:10.7554/eLife.88551

46. Sadakane O, Masamizu Y, Watakabe A, et al. Long-Term Two-Photon Calcium Imaging of Neuronal Populations with Subcellular Resolution in Adult Non-human Primates. Cell Rep. 2015;13(9):1989–1999. doi:10.1016/j.celrep.2015.10.050

47. Jendritza P, Klein FJ, Fries P. Multi-area recordings and optogenetics in the awake, behaving marmoset. Nat Commun. 2023;14(1):577. doi:10.1038/s41467-023-36217-5

48. Whishaw IQ, Gorny B. Arpeggio and fractionated digit movements used in prehension by rats. Behav Brain Res. 1994;60(1):15–24. doi:10.1016/0166-4328(94)90058-2

49. Bishop A. Control of the hand in lower primates. Ann N Y Acad Sci. 1962;102(2):316–337. doi:10.1111/j.1749-6632.1962.tb13649.x

50. Fox DM, Mundinano I, Bourne JA. Prehensile kinematics of the marmoset monkey: Implications for the evolution of visually-guided behaviors. J Comp Neurol. 2019;527(9):1495–1507. doi:10.1002/cne.24639

51. Takemi M, Kondo T, Yoshino-Saito K, et al. Three-dimensional motion analysis of arm-reaching movements in healthy and hemispinalized common marmosets. Behav Brain Res. 2014;275:259–268. doi:10.1016/j.bbr.2014.09.020

52. Fisk J, Lackner JR, DiZio P. Gravitoinertial force level influences arm movement control. J Neurophysiol. 1993;69(2):504–511. doi:10.1152/jn.1993.69.2.504

53. Gaveau J, Papaxanthis C. The Temporal Structure of Vertical Arm Movements. Choi DS, ed. PLoS ONE. 2011;6(7):e22045. doi:10.1371/journal.pone.0022045

54. Virji-Babul N, Cooke JD, Brown SH. Effects of gravitational forces on single joint arm movements in humans. Exp Brain Res. 1994;99(2). doi:10.1007/BF00239600

55. Papaxanthis C, Pozzo T, Stapley P. Effects of movement direction upon kinematic characteristics of vertical arm pointing movements in man. Neurosci Lett. 1998;253(2):103–106. doi:10.1016/S0304-3940(98)00604-1

56. Papaxanthis C, Pozzo T, Schieppati M. Trajectories of arm pointing movements on the sagittal plane vary with both direction and speed. Exp Brain Res. 2003;148(4):498–503. doi:10.1007/s00221-002-1327-y

57. Braccini SN, Caine NG. Hand preference predicts reactions to novel foods and predators in marmosets (Callithrix geoffroyi). J Comp Psychol. 2009;123(1):18–25. doi:10.1037/a0013089

58. Hook MA, Rogers LJ. Visuospatial reaching preferences of common marmosets (Callithrix jacchus): An assessment of individual biases across a variety of tasks. J Comp Psychol. 2008;122(1):41–51. doi:10.1037/0735-7036.122.1.41

59. Hashimoto T, Yamazaki Y, Iriki A. Hand preference depends on posture in common marmosets. Behav Brain Res. 2013;248:144–150. doi:10.1016/j.bbr.2013.04.001

60. Westergaard GC, Kuhn HE, Suomi SJ. Bipedal Posture and Hand Preference in Humans and Other Primates. J Comp Psychol. 1998;112(1):55–64. doi:10.1037/0735-7036.112.1.55

61. Cameron R, Rogers LJ. Hand preference of the common marmoset (Callithrix jacchus): Problem solving and responses in a novel setting. J Comp Psychol. 1999;113(2):149–157. doi:10.1037/0735-7036.113.2.149

62. Denenberg VH. Hemispheric laterality in animals and the effects of early experience. Behav Brain Sci. 1981;4(1):1–21. doi:10.1017/S0140525X00007330

63. Glick SD, Ross DA. Lateralization of function in the rat brain. Trends Neurosci. 1981;4:196–199. doi:10.1016/0166-2236(81)90063-1

64. Manns M, Basbasse YE, Freund N, Ocklenburg S. Paw preferences in mice and rats: Meta-analysis. Neurosci Biobehav Rev. 2021;127:593–606. doi:10.1016/j.neubiorev.2021.05.011

65. Gazzaniga MS. Principles of human brain organization derived from split-brain studies. Neuron. 1995;14(2):217–228. doi:10.1016/0896-6273(95)90280-5

66. Gotts SJ, Jo HJ, Wallace GL, Saad ZS, Cox RW, Martin A. Two distinct forms of functional lateralization in the human brain. Proc Natl Acad Sci. 2013;110(36). doi:10.1073/pnas.1302581110

67. Karolis VR, Corbetta M, Thiebaut De Schotten M. The architecture of functional lateralisation and its relationship to callosal connectivity in the human brain. Nat Commun. 2019;10(1):1417. doi:10.1038/s41467-019-09344-1

68. Oldfield RC. The assessment and analysis of handedness: The Edinburgh inventory. Neuropsychologia. 1971;9(1):97–113. doi:10.1016/0028-3932(71)90067-4

69. Papadatou-Pastou M, Ntolka E, Schmitz J, et al. Human handedness: A meta-analysis. Psychol Bull. 2020;146(6):481–524. doi:10.1037/bul0000229

70. Haaland KY, Delaney HD. Motor deficits after left or right hemisphere damage due to stroke or tumor. Neuropsychologia. 1981;19(1):17–27. doi:10.1016/0028-3932(81)90040-3

71. Haaland KY, Harrington D. The role of the hemispheres in closed loop movements. Brain Cogn. 1989;9(2):158–180. doi:10.1016/0278-2626(89)90027-4

72. Haaland KY, Harrington DL. Hemispheric asymmetry of movement. Curr Opin Neurobiol. 1996;6(6):796–800. doi:10.1016/S0959-4388(96)80030-4

73. Mutha PK, Haaland KY, Sainburg RL. The Effects of Brain Lateralization on Motor Control and Adaptation. J Mot Behav. 2012;44(6):455–469. doi:10.1080/00222895.2012.747482

74. Bagesteiro LB, Sainburg RL. Handedness: Dominant Arm Advantages in Control of Limb Dynamics. J Neurophysiol. 2002;88(5):2408–2421. doi:10.1152/jn.00901.2001

75. Bagesteiro LB, Sainburg RL. Nondominant Arm Advantages in Load Compensation During Rapid Elbow Joint Movements. J Neurophysiol. 2003;90(3):1503–1513. doi:10.1152/jn.00189.2003

76. Sainburg R. Evidence for a dynamic-dominance hypothesis of handedness. Exp Brain Res. 2002;142(2):241–258. doi:10.1007/s00221-001-0913-8

77. Sainburg RL, Kalakanis D. Differences in Control of Limb Dynamics During Dominant and Nondominant Arm Reaching. J Neurophysiol. 2000;83(5):2661–2675. doi:10.1152/jn.2000.83.5.2661

78. Sainburg RL, Schaefer SY. Interlimb Differences in Control of Movement Extent. J Neurophysiol. 2004;92(3):1374–1383. doi:10.1152/jn.00181.2004

79. Sainburg RL, Wang J. Interlimb transfer of visuomotor rotations: independence of direction and final position information. Exp Brain Res. 2002;145(4):437–447. doi:10.1007/s00221-002-1140-7

80. Schaefer SY, Haaland KY, Sainburg RL. Hemispheric specialization and functional impact of ipsilesional deficits in movement coordination and accuracy. Neuropsychologia. 2009;47(13):2953–2966. doi:10.1016/j.neuropsychologia.2009.06.025

81. Wang J, Sainburg RL. Interlimb transfer of visuomotor rotations depends on handedness. Exp Brain Res. 2006;175(2):223–230. doi:10.1007/s00221-006-0543-2

82. Przybyla A, Good DC, Sainburg RL. Dynamic dominance varies with handedness: reduced interlimb asymmetries in left-handers. Exp Brain Res. 2012;216(3):419–431. doi:10.1007/s00221-011-2946-y

83. Cabeza R. Hemispheric asymmetry reduction in older adults: The HAROLD model. Psychol Aging. 2002;17(1):85–100. doi:10.1037/0882-7974.17.1.85

84. Przybyla A, Haaland KY, Bagesteiro LB, Sainburg RL. Motor asymmetry reduction in older adults. Neurosci Lett. 2011;489(2):99–104. doi:10.1016/j.neulet.2010.11.074

85. Raw RK, Wilkie RM, Culmer PR, Mon-Williams M. Reduced motor asymmetry in older adults when manually tracing paths. Exp Brain Res. 2012;217(1):35–41. doi:10.1007/s00221-011-2971-x

86. Poirier G, Papaxanthis C, Lebigre M, et al. Aging decreases the lateralization of gravity-related effort minimization during vertical arm movements. Published online October 28, 2021. doi:10.1101/2021.10.26.465988

